# Automated detection of bacterial growth on 96-well plates for high-throughput drug susceptibility testing of *Mycobacterium tuberculosis*

**DOI:** 10.1101/229427

**Authors:** Philip W. Fowler, Ana Luíza Gibertoni Cruz, Sarah J. Hoosdally, Lisa Jarrett, Emanuele Borroni, Matteo Chiacchiaretta, Priti Rathod, Timothy M. Walker, Esther Robinson, Timothy E. A. Peto, Daniela Maria Cirillo, E. Grace Smith, Derrick W. Crook

## Abstract

*M. tuberculosis* grows slowly and is challenging to work with experimentally compared with many other bacteria. Although microtitre plates have the potential to enable high-throughput phenotypic testing of *M. tuberculosis*, they can be difficult to read and interpret. Here we present a software package, the Automated Mycobacterial Growth Detection Algorithm (`AMyGDA`), that measures how much *M. tuberculosis* is growing in each well of a 96-well microtitre plate. The plate used here has serial dilutions of 14 anti-tuberculosis drugs, thereby permitting the minimum inhibitory concentrations (MICs) to be elucidated. The two participating laboratories each inoculated ten 96-well plates with the standard H37Rv reference strain and, after two weeks incubation, measured the MICs for all 14 drugs on each plate and took a photograph. By analysing the images, we demonstrate that `AMyGDA` is reproducible, and that the MICs measured are comparable to those measured by a laboratory scientist. `AMyGDA` software will be used by the Comprehensive Resistance Prediction for Tuberculosis: an International Consortium (CRyPTIC) to measure the drug susceptibility profile of a large number (> 30,000) of samples of *M. tuberculosis* from patients over the next few years.

---

Tuberculosis (TB) kills more people globally than any other infectious disease [1, 2]. In 2016 the World Health Organisation (WHO) estimated that only 22% of the 600,000 people who required treatment for multi-drug resistant tuberculosis (MDR-TB) were diagnosed and received appropriate therapy [1]. In order to control the epidemic, priority should be given to the fast detection of MDR-TB cases and the identification of a proper and effective therapeutic regimen [3], which is currently done by culture-based drug-susceptibility testing (DST).

Existing liquid and solid media culture-based DST methods require significant infrastructure and highly-trained laboratory scientists and, due to the inherent slow-growth rate of *M. tuberculosis*, take at least 4-5 weeks to return a result to the clinician [4]. Although genotypic assays such as the Cepheid Xpert MTB/RIF [5] have been developed for *M. tuberculosis* (MTB) that overcome some of these challenges, no single molecular test exists that determines the effectiveness of a large number of anti-TB compounds simultaneously. One solution is to sequence the whole genome of each patient sample of *Mycobacterium tuberculosis* and infer the effectiveness, or otherwise, of a large number of drugs from the presence of genetic variants known to confer resistance [6, 7]. Using whole-genome sequencing (WGS) in this way has been shown to not only be much faster, but is already cheaper than the existing culture-based methods [4]. It critically depends, however, on a comprehensive and accurate catalogue that relates genetic variants to their effect on different anti-TB compounds. Whilst existing catalogues can reasonably accurately predict the effect of specific genetic mutations on the first-line anti-TB drugs (isoniazid, rifampicin, pyrazinamide, ethambutol) [8], more work is required to construct a catalogue allowing the effect of mutations on second-line, repurposed and novel compounds to be inferred, as well as improving the existing knowledge of first-line drugs. A comprehensive and accurate catalogue of genetic variants is vital for a WGS-based clinical microbiology to be able to accurately recommend treatments for MDR-TB [7].

The Comprehensive Resistance Prediction for Tuberculosis: an International Consortium [9] (CRyPTIC) is collecting over 30,000 samples of MTB world-wide over the next three years with the objective of identifying the majority of genetic variation in MTB responsible for antibiotic resistance in a large number of second-line, repurposed and novel anti-TB compounds, as well as the first-line compounds used in the standard regimen. The large number of samples involved make it too challenging and expensive to use traditional cultured-based methods. Microtitre plates are an attractive option for high-throughput culture-based DST, as they enable testing of a large number of strains in the presence of different anti-MTB drugs at a range of concentrations. One example is the Thermo Fisher Sensititre™ *M. tuberculosis* MIC Plate (MYCOTBI), which assays the minimum inhibitory concentration (MIC) for 12 anti-MTB drugs. Following inoculation of a cultured isolate, a plate is incubated for two weeks, and the presence or absence of growth in each well, and hence the MIC, is then manually assessed by a trained laboratory scientist. The accompanying Sensititre™ Vizion™ Digital MIC viewing system is designed to help with this process. CRyPTIC has developed a variant of the standard MYCOTBI plate, called the UKMYC5 plate, containing 14 different anti-MTB drugs, that includes two repurposed compounds (linezolid and clofazimine) and two new compounds (bedaquiline and delamanid, Fig. 1(a) & S1). The drugs are present at a minimum of 5 and a maximum of 8 doubling dilutions each [10]. Each *M. tuberculosis* patient sample collected by the CRyPTIC project will have its drug susceptibility profile determined using this plate. Since the main goal of the project is to combine the genetic and phenotypic data and statistically infer the effect of specific genetic variants, it is essential that all errors and biases in the data are minimised. Any system based on visual assessment by a trained laboratory staff member is, however, subject to some degree of variability between operators. Automated reading, using specifically-designed computer software, offers the promise of greater consistency but of course may not be as accurate, given a human’s superior ability to recognise and discriminate visual patterns.

**Figure 1:**
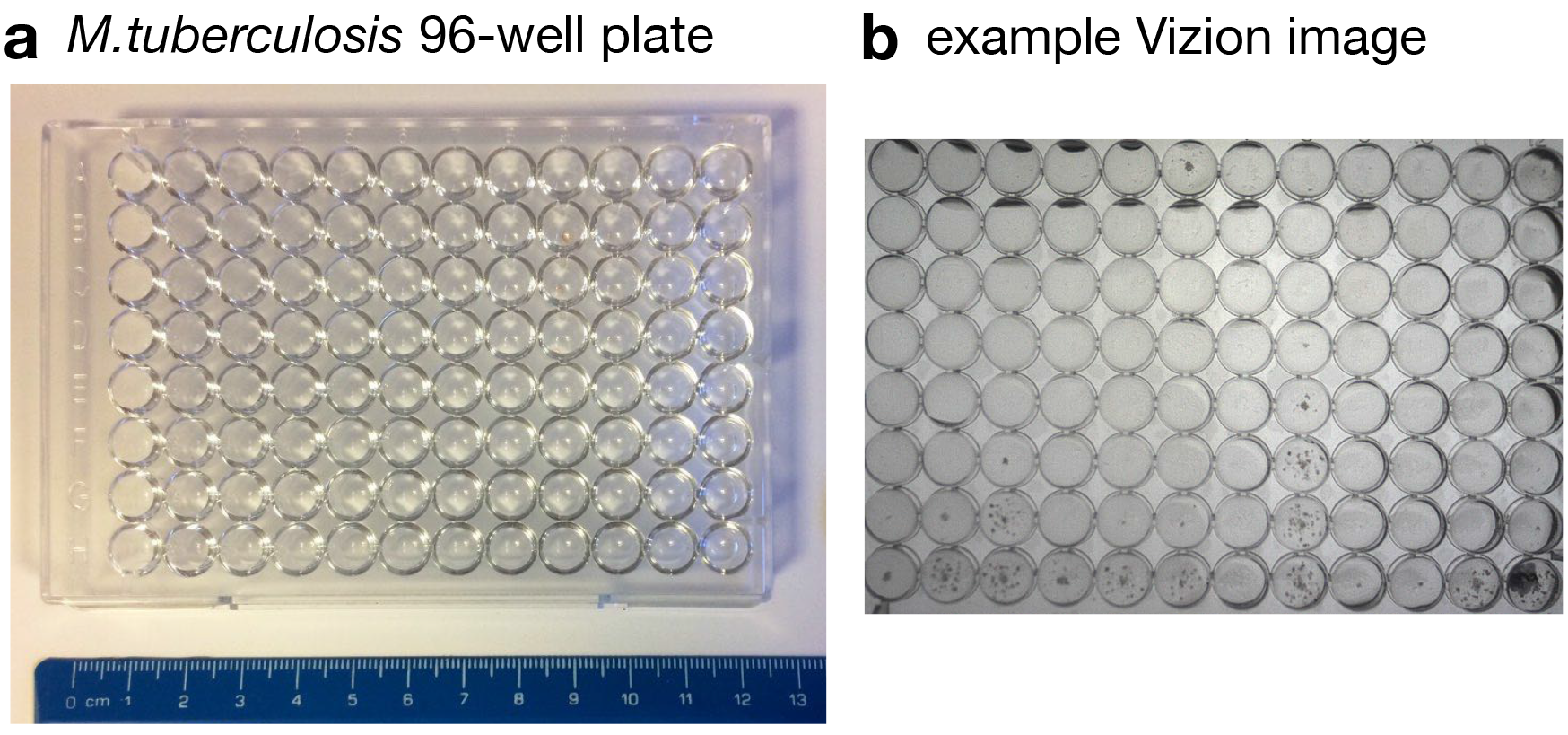
The 96-well plate used for *M. tuberculosis* (MTB) drug susceptibility testing. (**a**) The 96 well plate contains 14 different anti-MTB drugs. Each drug is present at 5-8 doubling dilutions and there are also two positive control wells which contain no drug. See Fig. S1 for the locations and concentrations of all 14 anti-MTB drugs. (**b**) Following inoculation with H37Rv and two weeks of incubation, an image is taken of each plate using a ThermoFisher Vizion™ Digital MIC viewing system. An example photograph is shown here. Specular growth can be seen in some wells. Raw images of all twenty plates can be found in Fig. S2

In this paper, we describe the design and parametrisation of software, the Automated Mycobacterial Growth Detection Algorithm (`AMyGDA`), that detects the growth of MTB on 96-well plates. To test its reproducibility and accuracy, we apply it to twenty UKMYC5 plates inoculated with the H37Rv ATCC 27294 MTB reference strain [11]. We intend deploying `AMyGDA` within the international CRyPTIC tuberculosis project to consistently determine MICs for the 14 anti-compounds on the UKUMYC5 plate for all >30,000 samples collected [10].

## METHODS

### Culturing

Each of the two laboratories received one screw-capped plastic cryovial with approximately 1000 *μ*l of mycobacterial suspension, and sub-cultured 200 *μ*l in a mycobacterial growth indicator tube (MGIT; Becton Dickinson). A second subculture step was then performed: 200 *μ*l of a well-mixed MGIT broth was inoculated on a solid Löwenstein-Jensen (LJ) medium, and incubated at 37 °C for 3-5 weeks. The MTB reference strain H37Rv, American Type Culture Collection 27294, obtained from the Biodefense and Emerging Infections Research Resources Repository (BEI Resources), USA was used throughout.

### Plate inoculation

2 to 5 mg of growth was harvested from LJ media after 3 to 5 weeks incubation, and re-suspended in 5 mL of saline solution, homogenized, and the supernatant density adjusted to a McFarland standard 0.5 suspension (~1.5 × 10^7^ CFU/mL). 100 *μ*l of this suspension was inoculated into a MGIT and 100 *μ*l dispensed from the MGIT into each of the plate’s 96 wells. Each plate was covered with an adhesive seal and transferred to aerobic incubation at 35-37°C

### Plate reading by the Vizion™ Digital MIC viewing system

Fourteen days after inoculation, laboratory staff read the plate using a Thermo Fisher Vizion™ Digital MIC reading system. MICs were recorded in a database. Vizion™ images were then generated according to the manufacturer’s guidance. Various combinations of Vizion™ lighting parameters – colour of background and level of illumination – were tested and, for the purposes of this study, the combination of ‘white background’ and ‘level 7’, respectively, was found to produce images with the highest contrast. Each image was manually cropped so it would fit on the Vizion™ Read window screen, with the result that final images had as similar dimensions as possible. Images were stored locally as lossless bitmap files (Fig. S2) by both laboratories before uploading to the database. In the Birmingham laboratory, the plate was then removed from the Vizion™ and a second laboratory scientist re-inserted the plate and took a second photograph, which was also uploaded to the database.

### Software

`AMyGDA` is written in object-oriented Python3. Each image is stored as a numpy array [12] and all image processing is done using the OpenCV2 Python API [13]. To allow for straightforward metadata storage we use the datreant module [14]. The algorithm assumes that each well is inoculated in the centre and mycobacteria will grow out radially from the centre, and that the images are 8-bit greyscale, allowing quantification of intensity of each pixel (range 0 to 255). Dark pixels (with low numerical values) were assumed to represent bacterial growth and light pixels to represent no growth. Growth was quantified according to the number of dark pixels per well. The `AMyGDA` software can be downloaded from http://fowlerlab.org/software/amygda.

### Filtering, image processing and growth detection

The raw images tend to be noisy, lacking in contrast and unevenly illuminated (Fig. 2(a), S2 & S3(a)). To correct for these problems a mean shift filter [15] is applied (Fig. 2(b), S3(b) & S4), followed by a contrast limited adaptive histogram equalization (CLAHE) filter (Fig. 2(c), S3D & S5). The positions of the wells are then detected using a Hough transform optimised for circles as implemented in OpenCV; this is applied iteratively until only 96 circles are detected in the image. Next, the histogram of pixel intensities in a central square region of each well is calculated and the proportion of bacterial growth inferred. If above a specified threshold, the well is labelled with a coloured square that also defines the region analysed for growth. This process, including the choice of parameters, is described in detail in the Supplementary Information. Finally, in addition to saving the MICs to disc, each image is annotated with the name and concentration of the drug in each well, providing a final composite image (Fig. 2(d), S12 & S13) that forms a complete, auditable record of the process.

**Figure 2:**
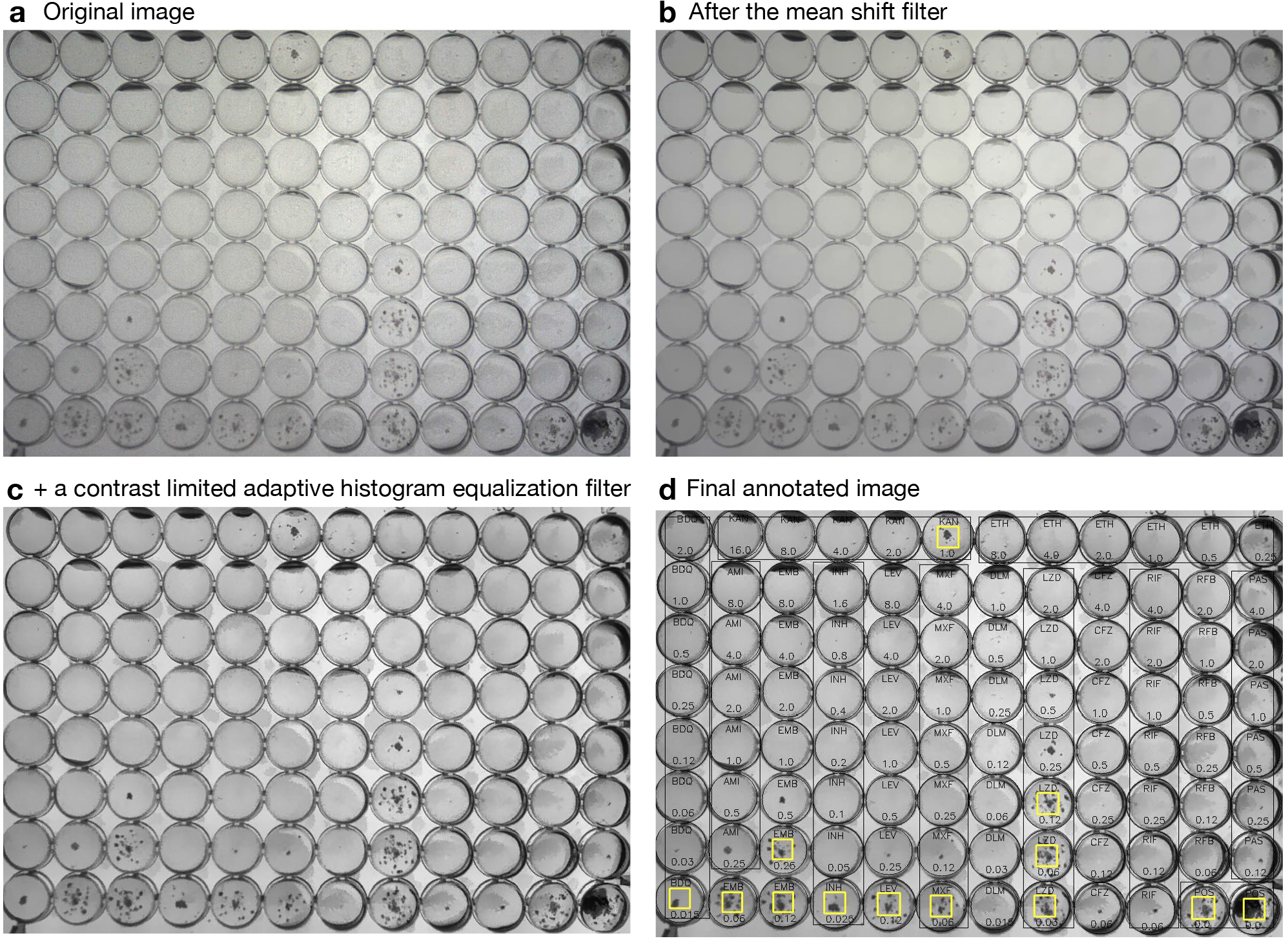
The AMyGDA software produces an annotated composite image with all detected growth labelled. (**a**) The original greyscale image is noisy and has poor contrast. To correct for this, (**b**) a mean shift filter and then (**c**) a contrast limited adaptive histogram equalization filter are applied. (**d**) The wells with detected bacterial growth are marked with a yellow square and each well is labelled with its drug and concentration and the estimated circumference is marked.

## RESULTS

### Study design

Two TB Reference laboratories (Birmingham, UK and Milan, Italy) received one vial of the *M. tuberculosis* ATCC27294 H37Rv reference strain. Each lab sub-cultured the isolate and inoculated ten UKMYC5 96-well microtitre plates. After two weeks of incubation, the MICs of the 14 anti-TB drugs on each plate were read by a trained laboratory scientist using a Vizion™ instrument, then a photograph of the plate was taken which was analysed at a later date by the `AMyGDA` software. To further test reproducibility, in one of the two laboratories a second scientist removed and re-inserted the plate into the Vizion and took a second photograph which was also analysed by the software. In total, 20 UKMYC5 plates were inoculated and incubated. Both `AMyGDA` and the laboratory scientist detected that *M. tuberculosis* appeared not to be growing in one of the two positive control wells of one plate: this plate was therefore excluded from analysis. In total, 19 plates were considered, resulting in 19 MICs for each of the 14 compounds, making 266 MICs in all.

### Sources of error

`AMyGDA` can wrongly report growth in a well – usually due to the presence of one of a range of image artefacts – and it can miss existing growth where only small and/or faint patches of growth are apparent. A trade-off between sensitivity and specificity is hence inevitable as these sources of error are inversely coupled. Artefacts which systematically affect a region of a plate or even the entire plate are more challenging to avoid or detect; these include shadows (Fig. 3(a)) and remnants of the original inoculation (‘sediment’, Fig. 3(b)). Artefacts which occur more randomly usually result in nonsensical growth patterns – e.g. the bacteria appear to grow at high, but not low, concentrations of antibiotic – which can be detected by the software and flagged for further investigation. These include air bubbles (Fig. 3(c)), condensation (Fig. 3(d)), contamination (Fig. 3(e)) and possible failure of the integrity of a plate, e.g. evaporation of inoculum during incubation due to inadequate plate sealing (Fig. 3(f)). For more information, including a detailed description of how the parameters were set within `AMyGDA` to maximise its sensitivity and specificity, please see the Supplementary Methods.

**Figure 3:**
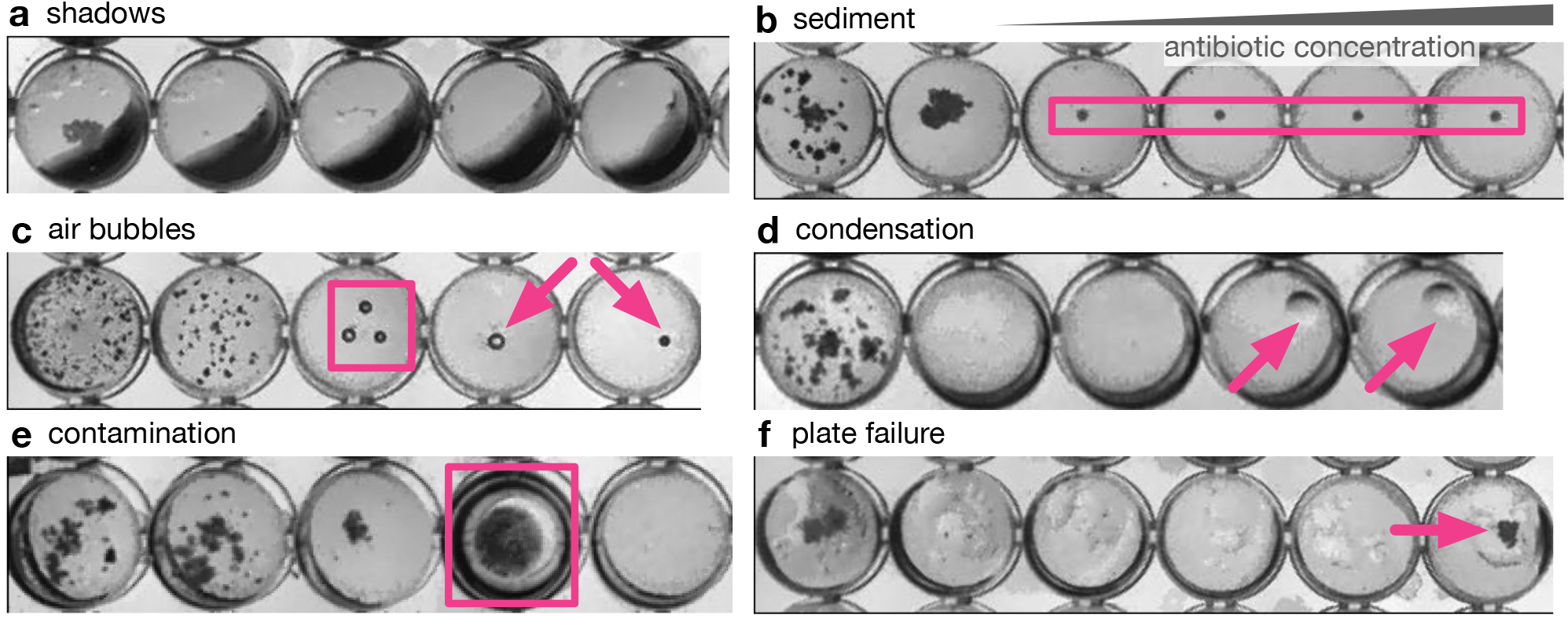
There are variety of artefacts that AMyGDA can mistake for growth, including (**a**) shadows, (**b**) sediment, (**c**) air bubbles, (**d**) condensation, (**e**) contamination and (**f**) possible failure of the plate integrity.

### Reproducibility of the software

As expected, `AMyGDA` produced identical results when applied ten times to the same image and is therefore reproducible. When applied to two different images of the same plate the amount of detected growth in each well varied slightly from one image to the other (Fig. 4(a)). To boost the number of wells in the analysis this comparison not used the two positive control wells but also a defined subset of drug-containing wells on the UKMYC5 plate for which the H37Rv reference strain tends to grow well (the lowest concentration wells for ethambutol, amikacin, isoniazid, levofloxacin, moxifloxacin and linezolid). Although a linear fit explains the data well, there are outliers and there is clearly some scatter as evidenced by a coefficient of determination of 0.93.

**Figure 4:**
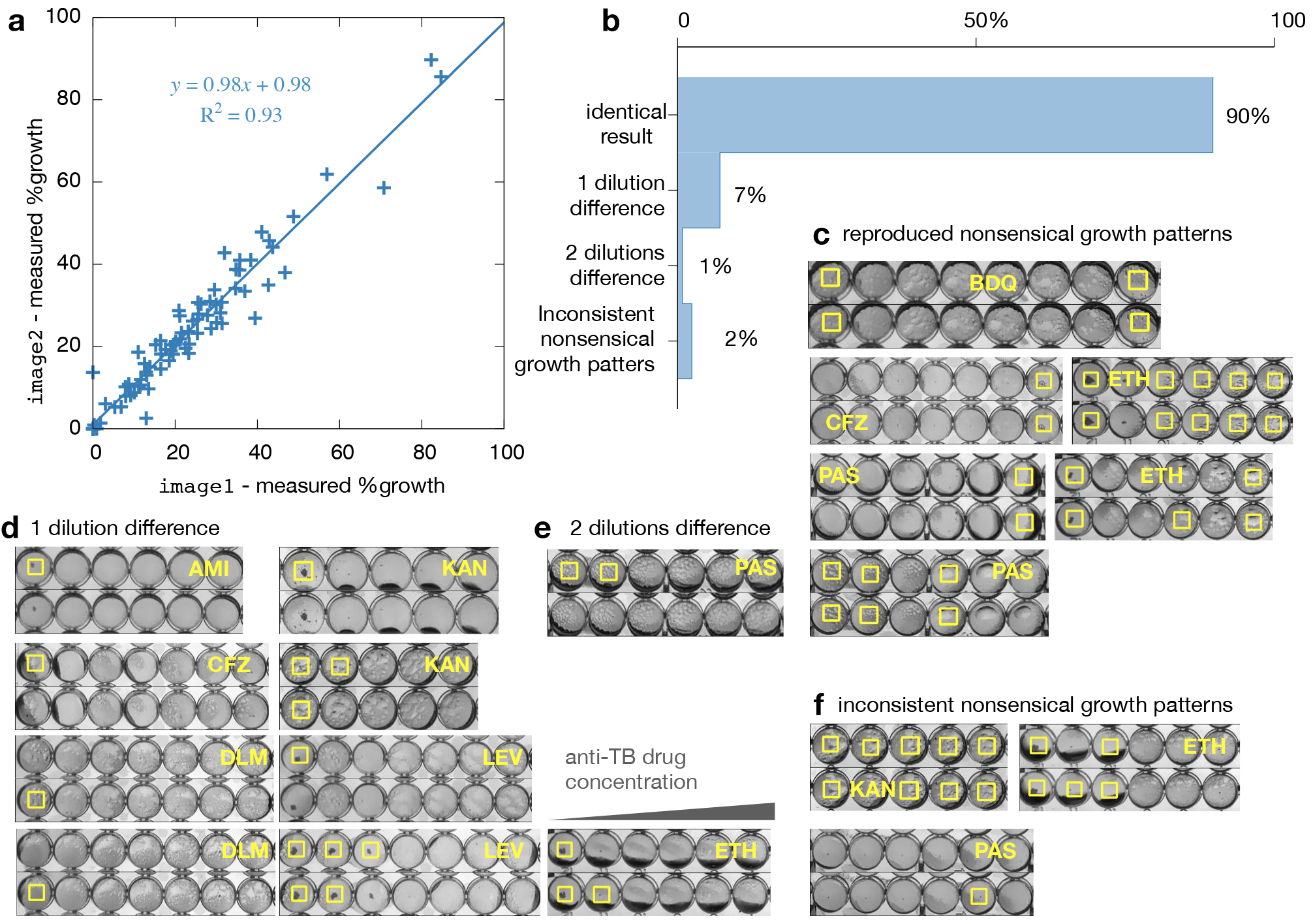
Applying the algorithm to the ten plates that were photographed twice (Fig. S14) measures the same MIC for 90% of wells. One of the ten replicates was excluded as the algorithm only detected growth in one of the two control wells. (**a**) The growth measured in a subset of wells on image1 and image2 is correlated. (**b**) The algorithm infers exactly the same MIC in 90% (113/126) of cases. This is comprised of 107 cases where the minimum inhibitory concentration is identical (examples not shown) and (**c**) six cases (5%) where there is a nonsensical growth pattern in both images. (**d**) There are nine cases (7%) where there is a single doubling dilution difference. (**e**) There is a single case where there are two doubling dilutions difference. (**f**) Finally, in three cases (2%) a nonsensical growth pattern is returned for one image, but not the other.

Due to the one plate being excluded, there were nine UKMYC5 plates with two images available for this analysis, making a total of 126 pairs of MIC measurements that can be compared. In 90% of cases (113/126) `AMyGDA` estimated an identical MIC (Fig. 4(b) & S14) when analysing the second image, including six cases where a nonsensical growth pattern was detected in both images, namely where growth was detected in higher but not in lower concentration wells (Fig. 4(c)). Of the remaining 10% (13/126), the MIC measured by `AMyGDA` from the second image was only 1 doubling dilution different in the majority (7%, 9/126) of cases (Fig. 4(c)). Examining the images in more detail shows that these cases were a mixture of wells which either had little growth or artefacts. One MIC measured from the second image was 2 doubling dilutions lower than the MIC measured from the first image (Fig. 4(d)), due to condensation in this antibiotic lane being incorrectly classified as growth in two consecutive wells in one image, but not in the other. The differences in the remaining 2% (3/126) of MICs pairs were when the algorithm detected a nonsensical growth pattern in one image, but not in the other.

There is no international standard for Mycobacterial antimicrobial susceptibility testing (AST), therefore there is no agreed threshold that any new AST method must clear before it can be considered for clinical use. We shall therefore apply with caution the standard defined for aerobic bacteria [16]. This states that the proportion of any readings within a doubling dilution of the mode must be ≥ 95%. On the basis of this small dataset, `AMyGDA` achieves 96.8% (122/126), which is promising but not yet conclusive.

### Validation against measurements made by a laboratory scientist

As mentioned above, prior to analysis by `AMyGDA`, all UKMYC5 plates were read by a trained laboratory scientist using the Vizion™ Digital MIC instrument, allowing us to compare the MICs obtained by both methods (Fig. 5(a)). There were no systematic differences in the MICs measured by both laboratories.

**Figure 5:**
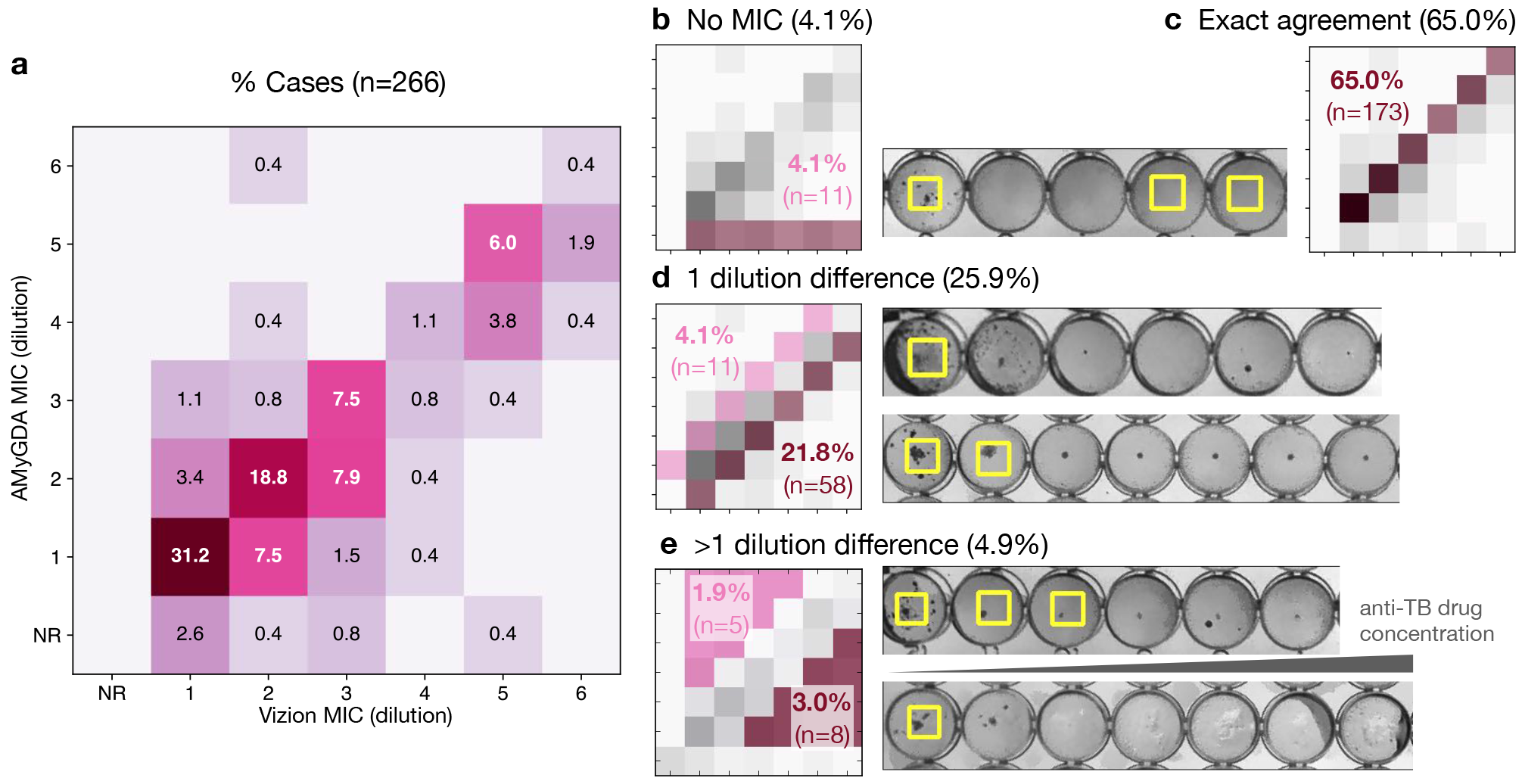
The results of the automated mycobacterial growth detection algorithm (AMyGDA) were validated by comparing to independent measurements made by laboratory scientists using the Vizion™ Digital MIC viewing instrument. (**a**) A heat map for all MIC doubling dilutions from all twenty test plates showing the concordance between the human-based measurement and the AMyGDA software. (**b**) Both measurements reject one plate (5%) which has no or unusual growth in one of the control wells. The AMyGDA software identifies nonsensical growth in a further 3.9% of cases. (**c**) Both methods infer the same MIC in 61.8% of cases with (**d**) a further 24.6% within ±1 doubling dilution. (**e**) In only 4.7% of cases do the methods disagree by more than a doubling dilution.

The `AMyGDA` software was unable to return an MIC in 11/266 of cases (4.1%) due to nonsensical growth patterns. These were mostly due to artefacts being incorrectly classified as bacterial growth (Fig. 5(b)). In all 11 cases the laboratory scientists were able to discern an MIC, demonstrating the superior ability of a human to identify and ignore artefacts. For 173/266 (65%) of cases, both methods gave exactly the same MIC (Fig. 5(c)). In 58/266 (21%) of cases `AMyGDA` reported an MIC one doubling dilution lower than the laboratory scientist. In contrast, in only 11/280 (4%) of cases did the software predict an MIC one doubling dilution higher than the scientist. This disparity is due to the relatively conservative setting of the growth classification parameters to minimise the detection of artefacts as described in the Supplement. Finally, for 13 MICs, the laboratory scientist and the `AMyGDA` software assigned MICs that were two or more dilutions different (Fig. 5(e)), usually because there was little growth in the wells.

There is no accepted threshold for accuracy due to the lack of a Mycobacterial AST standard, so again we shall tentatively apply the criteria any new AST method for aerobic bacteria must satisfy when its results are compared to those of an accepted AST method [16]: the new method must be in essential agreement (EA) in ≥ 90% of cases. Essential agreement is defined as the two MICs lying within a single doubling dilution of one another. The `AMyGDA` software has an essential agreement of 90.9% (242/266) with the Vizion readings. This should not be taken as evidence that the software is therefore sufficiently accurate to be deployed clinically, however, since there are several problems with this analysis that we will discuss shortly.

## DISCUSSION

We have shown how `AMyGDA`, a Python package, can detect and measure growth of *M. tuberculosis* in images of a 96-well microtitre plate, each having been inoculated and incubated with the H37Rv *M. tuberculosis* strain. Here we used a variant of the standard Thermo Fisher Sensititre™ MYCOTB 96-well plates, UKMYC5, that has been designed by the international CRyPTIC tuberculosis consortium. Since UKMYC5 contains 14 different anti-TB compounds, each forming a doubling dilution series, the software therefore can determine the minimum inhibitory concentration for each of the 14 drugs on the UKMYC5 plate.`AMyGDA` can be downloaded from http://fowlerlab.org/software/amygda.

Ideally one applies an external set of success criteria when assessing the effectiveness of new software. As there are no antimicrobial susceptibility testing (AST) standards for Mycobacteria, we have tentatively applied an international AST standard for aerobic bacteria [16]. This is a stringent test since any new AST test must pass the appropriate standard before it can be considered for use clinically. The results indicate that `AMyGDA` is *potentially* sufficiently reproducible and accurate to pass the thresholds defined in the standard and therefore we conclude that `AMyGDA` is suitable for use in a high-throughput research setting. It would be inappropriate, however, to conclude that `AMyGDA` is suitable for deploying in clinical microbiology laboratories. For that to be determined, a much larger and more comprehensive validation study using clinical isolates will be required to more systematically evaluate the accuracy and reproducibility of the `AMyGDA` software.

The CRyPTIC project has therefore adopted `AMyGDA` primarily to assist in the quality control and identification of discrepancies. Since photographs will be taken of all plates after two weeks incubation, the intention is to retrospectively analyse all these images and, by merging the measurements taken by the expert, identify and investigate plates where there is insufficient agreement between the MICs measured by human and computer. In this way we will identify measurement errors and so minimise the biases and errors in the large phenotype dataset that the CRyPTIC project is collecting. This large dataset will also be sufficiently large enough to conclusively demonstrate, given a Mycobacterial AST standard, that `AMyGDA` is sufficiently reproducible and accurate for clinical use.

Since `AMyGDA` measures the amount of growth in each well, it also potentially opens up new research questions. For example, as the concentration of drug is increased, how does the apparent growth of *M. tuberculosis* change from well to well? Or, more ambitiously, by linking to the genomic data that is being collected, can we identify strains or mutations that grow quickly or better (at least on the UKMYC5 plate)? These and other questions will be the focus of future studies.

An additional use for this and similar automated plate-reading software could be for high-throughput phenotypic screening for *M. tuberculosis* drug discovery. Unlike a human, the software is quantitative and so could, given enough samples, detect small changes in the rate of growth due to mild inhibition, enabling the use of microtitre plates to identify potential leads for novel antibiotic compounds by high-throughput phenotypic screening of a compound- or fragment-library which otherwise would be missed.

Conventionally culture-based methods are read after a fixed time period: here two weeks following inoculation. We observed, however, significant variation in the amount of growth between plates even though these were all inoculated with the same strain, suggesting variability in inoculation has a potentially significant effect. This is likely to be accentuated further when different strains are compared. Software, like `AMyGDA`, could instead prospectively monitor the growth in the control wells during the incubation period, allowing a plate to only read once there is ‘sufficient’ growth (‘read-when-ripe’), further reducing the time between sample collection and result.

There are a number of limitations to our study in addition to the ones mentioned above. The Vizion™ was not designed to capture photographs that could be processed by software and hence it has uneven illumination and the digital camera is, by today’s standards, of low resolution. An alternative, cheaper way of capturing images with more diffuse illumination and a higher resolution camera would help considerably. Secondly, whilst we have minimised the false detection of artefacts, they remain a problem, especially those that tend to systematically afflict whole regions of a plate. Improved experimental protocols may help avoid some of these (e.g. air bubbles) and a re-design of the plate layout would help with others (e.g. shadows).

In conclusion, combining computer software, such as `AMyGDA`, with microtitre plate-based MIC measurement can facilitate high-throughput culture-based drug susceptibility testing for tuberculosis.

## ACKNOWLEDGEMENTS

We are grateful to members of the Oxford Doctoral Training Centre 2016/7 cohort for algorithmic suggestions. The research was funded by the National Institute for Health Research (NIHR) Oxford Biomedical Research Centre(BRC); the CRyPTIC consortium which is funded by a Wellcome Trust/Newton Fund-MRC Collaborative Award [200205/Z/15/Z] and the Bill & Melinda Gates Foundation Trust [OPP1133541]. T.E.A.P. and D.W.C. are NIHR Senior Investigators. T.M.W. is an NIHR Academic Clinical Lecturer. The views expressed are those of the author(s) and not necessarily those of the NHS, the NIHR or the Department of Health.

## References

[1] World Health Organisation (2017) Global Tuberculosis Report. Technical report.

[2] Fogel N (2015) Tuberculosis 95:527–531.

[3] World Health Organisation (2011) Towards universal access to diagnosis and treatment of multidrug-resistant and extensively drug-resistant tuberculosis by 2015. Technical report.

[4] Pankhurst LJ, del Ojo Elias C, Votintseva AA, Walker TM, Cole K, Davies J, Fermont JM, Gascoyne-Binzi DM, Kohl TA, Kong C, Lemaitre N, Niemann S, Paul J, Rogers TR, Roycroft E, Smith EG, Supply P, Tang P, Wilcox MH, Wordsworth S, Wyllie D, Xu L, Crook DW (2016) Lancet Resp Med 4:49–58.

[5] Boehme CC, Nabeta P, Hillemann D, Nicol MP, Shenai S, Krapp F, Allen J, Tahirli R, Blakemore R, Rustomjee R, Milovic A, Jones M, O’Brien SM, Persing DH, Ruesch-Gerdes S, Gotuzzo E, Rodrigues C, Alland D, Perkins MD (2010) New Eng J Med 363:1005–1015.

[6] Didelot X, Bowden R, Wilson DJ, Peto TEA, Crook DW (2012) Nat Rev Genetics 13:601–12.

[7] Ellington M, Ekelund O, Aarestrup F, Canton R, Doumith M, Giske C, Grundman H, Hasman H, Holden M, Hopkins K, Iredell J, Kahlmeter G, Köser C, MacGowan A, Mevius D, Mulvey M, Naas T, Peto T, Rolain JM, Samuelsen Ø,Woodford N (2017) Clinical Microbiology and Infection 23:2–22.

[8] Walker TM, Kohl TA, Omar SV, Hedge J, Del Ojo Elias C, Bradley P, Iqbal Z, Feuerriegel S, Niehaus KE, Wilson DJ, Clifton DA, Kapatai G, Ip CLC, Bowden R, Drobniewski FA, Allix-Béguec C, Gaudin C, Parkhill J, Diel R, Supply P, Crook DW, Smith EG, Walker AS, Ismail N, Niemann S, Peto TEA, Modernizing Medical Microbiology (MMM) Informatics Group (2015) Lancet Infec Disease 15:1193–202.

[9] Comprehensive Resistance Prediction for Tuberculosis: an International Consortium (2017). http://www.crypticproject.org.

[10] Rancoita PMV, Cugnata F, Luíza A, Cruz G, Hoosdally SJ, Walker TM, Grazian C, Davies TJ, Peto TEA, Crook DW, Fowler PW, Cirillo DM (2018) biorXiv preprint Submitted.

[11] Kubica GP, Kim TH, Dunbar FP (1972) Int J System Bacteriol 22:99–106.

[12] Van Der Walt S, Colbert SC, Varoquaux G (2011) Comp Sci Eng 13:22–30.

[13] Bradski G, Kaehler A (2008) Learning OpenCV. O’Reilly Media, Inc, first edition.

[14] Dotson DL, Seyler SL, Linke M, Gowers RJ, Beckstein O (2016) In Proc 15th Python Sci Conf, edited by S Benthall, S Rostrup, 51–56.

[15] Comaniciu D, Meer P (2002) IEEE Trans Pattern Analysis Mach Intel 24:603–619.

[16] International Organization for Standardization (2007) ISO 20776-2: Clinical laboratory testing and in vitro diagnostic test systems. Technical report, International Standards Organization.

